# Transposable elements in a cold-tolerant fly species, *Drosophila montana*: a link to adaptation to the harsh cold environments

**DOI:** 10.1101/2024.04.17.589934

**Authors:** Mohadeseh S. Tahami, Carlos Vargas-Chavez, Noora Poikela, Marta Coronado-Zamora, Josefa González, Maaria Kankare

## Abstract

**Background:** Substantial discoveries during the past century have revealed that transposable elements (TEs) can play a crucial role in genome evolution by affecting gene expression and inducing genetic rearrangements, among other molecular and structural effects. Yet, our knowledge on the role of TEs in adaptation to extreme climates is still at its infancy. The availability of long-read sequencing has opened up the possibility to identify and study potential functional effects of TEs with higher precision. In this work, we used *Drosophila montana* as a model for cold-adapted organisms to study the association between TEs and adaptation to harsh climates.

**Results:** Using the PacBio long-read sequencing technique, we *de novo* identified and manually curated TE sequences in five *Drosophila montana* genomes from eco-geographical distinct populations. We identified 489 new TE consensus sequences which represented 92% of the total TE consensus in *D. montana*. Overall, 11-13% of the *D. montana* genome is occupied by TEs, which as expected are non-randomly distributed across the genome. We identified five potentially active TE families, most of them from the retrotransposon class of TEs. Additionally, we found TEs present in the five analyzed genomes that were located nearby previously identified cold tolerant genes. Some of these TEs contain promoter elements and transcription binding sites. Finally, we detected TEs nearby fixed and polymorphic inversion breakpoints.

**Conclusions:** Our research revealed a significant number of newly identified TE consensus sequences in the genome of *D. montana*, suggesting that non-model species should be studied to get a comprehensive view of the TE repertoire in Drosophila species and beyond. Genome annotations with the new *D. montana* library allowed us to identify TEs located nearby cold tolerant genes, and present at high population frequencies, that contain regulatory regions and are thus good candidates to play a role in *D. montana* cold stress response. Finally, our annotations also allow us to identify for the first time TEs present in the breakpoints of three *D. montana* inversions.

## Background

Environmental stress is one of the key traits affecting species survival and distribution, especially in northern areas where temperature fluctuations are less predictable and extreme weather events are becoming more frequent because of climate change [1,2]. Recent studies have indicated that species may adapt through various interplays between different genetic elements in the organism’s genome [3–7]. One of the most interesting genetic elements connected to adaptation are transposable elements (TEs). TEs are repetitive sequences with the ability to replicate themselves independently of the genome and change their position thereby generating diverse mutations. They represent a significant portion of the genome in many species, for example, over two thirds of the human genome [8] and 90% of the wheat genome [9] are TEs. In insects, TE content also contributes significantly to genome size variation [10], for example, in *Drosophila* flies, TE content can vary between 3% and 30% [11]. The activity and the abundance of these highly repetitive sequences in the genome suggests that TEs could be important factors in the evolution of many species.

TEs can impact species adaptation to environmental changes. For example, they are known to be activated under stress [12] and can affect the species response to environmental stressors such as heat shock [13], cold [14], pesticides [15], insecticides [16] and oxidative stress [17] thus creating a tolerance against these stressors. TEs can have diverse effects on the host’s genome based on the location they have been inserted and the regulatory sequences they contain, *e.g.* they can create insertional mutations, up-or downregulate nearby genes, or create new gene variants by introducing new exons, new stop codons, or alternative splice sites [18–20]. In some cases, one of these newly generated alleles may become adaptive in response to environmental stress. This so-called ‘adaptive TE’ will be co-opted and thus tolerated by natural selection. The case of the co-opted *FBti0019985* transposon located in the promoter of the *Lime* gene of *D. melanogaster* is an example in which the adaptive TE affects gene expression associated with cold-stress and immune tolerance [21,22]. In the case of immune tolerance, the authors show that *FBti0019985* insertion is adding functional transcription factor binding sites (TFBS) for transcription factors involved in immune response [22]. If the stress conditions persist, co-opted TEs might become fixed in the population [23].

Another way that TEs can participate in environmental adaptation is by inducing ectopic recombination leading to structural variations such as inversions [24,25]. One such case is reported for a Galileo transposon that has generated three polymorphic inversions in *Drosophila buzzatii* through ectopic recombination [26]. Inversions in turn can play a role in species adaptation and speciation in many ways [reviewed in e.g. 27–29]. Their main evolutionary significance is that they can reduce recombination between favorable combinations of alleles while protecting sets of locally adapted genes, thereby promoting ecological divergence, and fostering reproductive isolation within species [30]. For example, inversions are considered to be prevailing drivers of population divergence in *D. virilis* species group [31]. Despite the abundance of both intraspecific polymorphic and interspecific fixed inversions in Diptera, studies on the role of TEs on chromosomal inversions are still scarce.

The northern malt fly, *Drosophila montana*, is a widely distributed insect species which belongs to the *Drosophila virilis* group [32] and is one of the most cold-tolerant *Drosophila* species [33,34]. These northern malt flies can spend up to 6 months in subzero temperatures and several populations have adapted to live in latitudes above the Arctic Circle in Northern Scandinavia. In lower latitudes, flies inhabit high elevations (about 3,000 m) e.g., in the Rocky Mountains in Colorado (USA). They also exist in the warmer coastal areas in Washington and Oregon (USA) and hence show a wide range of eco-climate adaptation [35]. Populations of *D. montana* flies are clustered into European, North American and Asian populations [36] and the current geographical pattern is likely the result of spreading North American populations into Eurasia through the Bering Strait between 450,000 and 1,750,000 years ago [35] resulting in its unique intraspecific diversification. In *D. montana*, adaptation to the extreme climate is modulated by several genes associated with cold tolerance [37–43]. These genes regulate a variety of functions from general cold resistance to cuticular and olfactory processes and photoperiodic diapause, enhancing the species overwintering ability [34,42]. It is also possible that some TEs may have a role in the regulation of the cold-related genes, leading to the species high resistance e.g. to the freezing temperature. Moreover, based on earlier polytene chromosome studies, *D. montana* carries several fixed and polymorphic inversions [44,45]. So far, one fixed inversion [46] (in comparison to *Drosophila flavomontana*), and two polymorphic inversions (Poikela et al., in prep.) has been characterized at the genomic level, but the link between TEs and inversions in *D. montana* has not been studied yet.

Here, we identified TEs in *D. montana* flies across its distribution range for the first time using long-read genome sequence data. To study the potential role of TEs in adaptation to northern environments, we first produced a comprehensive manually curated TE library from North American, North European and Asian populations of *D. montana*. Next, we investigated the abundance, density, distribution and activity of TEs throughout these genomes. Finally, we studied the association between TEs and chromosomal inversions, as well as the link between TEs and a selected set of cold tolerant candidate genes found in previous studies to identify possible adaptive TEs. We addressed the following questions: (i) what is the genome-wide TE profile in *D. montana* and does it differ from that of other *Drosophila* species? (ii) Can we detect potentially active TEs in the *D. montana*’s genome? (iii) Can we find any evidence for the role of TEs in adaptation to the northern environments? and (iv) are there TE insertions located nearby inversion breakpoints?

## Methods

### Sample collections

*Drosophila montana* flies were collected from 13 locations in North America (NA), North Europe (NE), and Far East Asia (FE) between 2013 and 2021 (Figure 1 and Supplementary Table S1). The collected females were brought to the fly laboratory of the University of Jyväskylä, Finland, and kept in constant light, 19 °C and ∼60% humidity. The females that had mated in nature were allowed to lay eggs in malt vials for several days. The emerged F1 progeny of each female were kept together to produce the next generation and to establish isofemale strains. After the establishment of isofemale strains, all wild-collected females and their F1 progeny, except those collected in FE, were stored in 70% EtOH at -20 °C.

**Figure 1.**
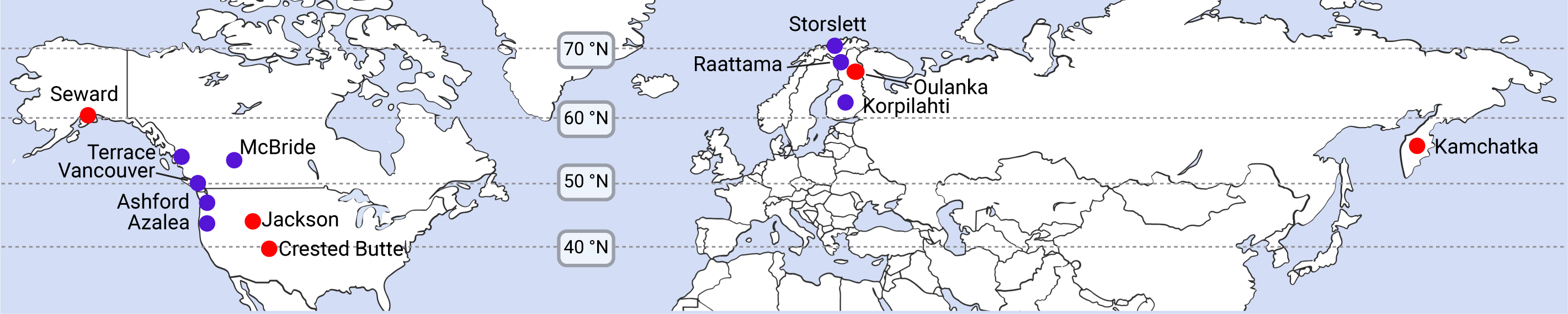
Sample locations of the *D. montana* populations analyzed in this study. The red circles indicate the location for the five genomes that were used to generate the original TE library (supplementary Table S2). The red and blue circles show the location of populations used in the Illumina sequencing (supplementary Table S3).

For PacBio (Pacific Biosciences) long-read sequencing, *D. montana* females were collected from five isofemale strains originating from five locations: three in NA, Seward (monSE13F37; Alaska, USA) and Jackson (monJX13F48; Wyoming, USA) from a previous study [46] and Crested Butte (mon34CC5; Colorado, USA) from the current study; one in NE, Oulanka (monOU13F149; Finland); and one in FE, Kamchatka (monKR1309; Russia) (Figure 1 and Supplementary Tables S1 & S2). Prior to collecting the flies for PacBio sequencing, isofemale strains were kept in the laboratory for ∼50 generations (Supplementary Table S2).

For Illumina short-read sequencing, we used the stored wild-collected females or their F1 daughters from 12 collection sites in NA and NE. Because the wild-collected females or their F1 daughters from FE were not stored at -20 °C, we performed Illumina sequencing on females collected from isofemale strains kept in the laboratory for ∼60 generations (Figure 1 and Supplementary Tables S1 & S3)

### Long-read sequencing

PacBio sequencing technique was used for the five *D. montana* isofemale strains. Except for Seward and Jackson [46], all samples were sequenced in this study. For all samples, DNA was extracted from a pool of 60 females. Details of samples, DNA extraction, and sequencing methods are given in Supplementary Table S2.

### Short-read sequencing

Illumina whole-genome sequencing was performed for 3-11 wild-collected or F1 females per population from NA and NE, and for 3 females collected from FE isofemale strains (Figure 1 and Supplementary Table S3). The majority of the fly samples (94/101) were sequenced for this study, but seven samples were obtained from Poikela et al. [46] (see Supplementary Table S3). DNA extractions were carried out for single females using cetyltrimethylammonium bromide (CTAB) solution with RNAse treatment, Phenol-Chloroform-Isoamyl alcohol (25:24:1) and Chloroform-Isoamyl alcohol (24:1) washing steps and ethanol precipitation at the University of Jyväskylä, Finland. Details of library preparation, sequencing technologies, locations, and years are given in Additional file 1, Supplementary Table S3. We generated on average 19-97X coverage per sample (Supplementary Table S3).

### *De novo* genome assemblies and scaffolding

We obtained *de novo* genome assemblies for *D. montana* Seward and Jackson from Poikela et al. [46] and constructed new *de novo* genome assemblies for *D. montana* Crested Butte, Kamchatka and Oulanka following the same protocol. In brief, we assembled the genomes using PacBio and the respective Illumina reads with *wtdbg2* pipeline v2.5 [47] and MaSuRCA hybrid assembler v3.3.9 [48], after which the assembly contiguity was improved using *quickmerge* v0.3 [49] and the assemblies were polished with the same Illumina reads using *Pilon* v1.23 [50]. Finally, uncollapsed heterozygous regions were removed using *purge_dups* v1.0.1 [51] and genomic contaminants were removed using *BlobTools* v1.1 [52]. We estimated the completeness of the assemblies using the BUSCO pipeline v5.1.2 based on the Diptera database “diptera_odb10” [53], which searches for the presence of 3,285 conserved single copy Diptera orthologues.

Chromosome-level genome assembly for *D. montana* Seward was also obtained from Poikela et al. [46]. This assembly was used to scaffold the rest of the genomes (Jackson, Crested Butte, Kamchatka and Oulanka) using a reference-genome-guided scaffolding tool *RagTag* v2.1.0 [54] which orients and orders the input contigs based on a reference using *minimap2* [55].

### Building a *de novo* TE library

The five *de novo* assembled genomes were used for *de novo* identification of TEs using the *TEdenovo* pipeline in the *REPET* package v3.0 [56]. In short, repeated regions were detected by self-alignment of the genomic chunks, the TE candidates were clustered using three methods: *Recon* v1.08, *Grouper* v2.27 and *Piler* v1.0. Consensus sequences were derived from multiple alignments in each cluster. The consensuses were classified based on structural features or homology match with the *Repbase* reference library v20.05 [57]and hidden Markov model (HMM) profiles. To annotate consensus TEs, each TE library was blasted against its host genome using three different tools *Blaster* v2.25, *RepeatMasker* v4.0.6, and *CENSOR* v4.1, integrated into *TEannot* pipeline [56]. To apply a statistical filtering, a randomized genomic chunk alignments was also applied in order to calculate the high-scoring segment pairs (HSP). The next steps filtered and combined HSPs by keeping only the ones having a score higher than the HSP threshold, removed spurious HSPs, and calculated identity percentages. We also kept and merged the short simple repeat (SSR) data by applying step 4 and 5 of *TEannot*. Then, consensus sequences with less than one full length copy throughout the genome were filtered out and the TEannot process was repeated on the updated library. For manual curation, the consensus sequences of each individual library were visualized in the *Integrative Genomics Viewer* (IGV) v2.8.0 [58] along with their structural features, conserved domains, nucleotide, and amino acid homology matches. We curated identified TEs based on Wicker’s features [59] and we followed several steps to reduce the false positive error in our TE library and to remove the redundancy. For more details on the manual curation process please refer to the Additional file 1, Supplementary Table S4 and Additional file 2. To validate our manual curation process, we run *MCHelper* on the final library [60].

### Heterochromatin identification

Because we were interested in TE insertions with potential functional impact, we identified heterochromatin regions to be removed from further TE analyses. After the manual TE curation, the final library was used to annotate TEs across each chromosome-level assembly using *RepeatMasker* v4.1.0 [61] with following parameters: -e ncbi -nolow -xsmall -gccalc. Since no prior information on the heterochromatin regions of *D. montana* genome was available, we plotted the total TE densities per 5 kb genomic bins. We considered the regions with the highest TE density to be heterochromatic regions. To view the TE density plot please refer to Additional file 3 Supplementary Figure S1 and for the table hetero-vs euchromatin coordinates, refer to Additional file 1, Supplementary Table S5. We identified these regions as those that have distinctive long stretches of peaks followed by a sharp decrease in TE density and all these regions were located at the ends of each chromosomal arm, as expected.

### Gene annotation

We used the Seward chromosome-level assembly as the reference for genome annotation. The curated TE library of *D. montana* was used to mask genomes using *RepeatMasker* v4.1.0. The soft-masked genome was annotated using *D. montana* RNAseq data (Illumina Truseq 150bp PE) [62]. RNA-seq reads were first trimmed using *fastp* v0.20.0 [63] and mapped against the soft-masked genome using *STAR* v2.7.10a [64]. Gene annotations were carried out with *braker* v2.1.6 [65] with default parameters, with RNAseq as evidence using *Augustus* v3.4.0 and *GeneMark-ET* v4.48 implemented pipelines [66–71]. Finally, the annotation was transferred to the other chromosome-level genomes using *Liftoff* v1.6.3 [72]. A range of 14,504 to 14,725 genes were transferred to each genome (Supplementary Table S6). The average error rate was calculated based on the total number of exons that were correctly transferred for each gene to be at 1.1%.

### SNP calling

To examine if SNP variation within TE regions can reflect the geographical variation between *D. montana* genomes across its distribution, Illumina reads corresponding to each *D. montana* genome assembly were used to extract nucleotide variants from the whole genome and from TE regions for comparison. Reads were filtered for adapters and bases below quality of 20 were trimmed at both ends using *fastp* v0.21.0. Trimmed reads were mapped to the Seward chromosome assembly (monSE13F37) using *BWA mem* v0.7.17 [73] with default parameter settings. PCR duplicates were removed from sorted bam files with *sambamba* v 0.7.0 [74]. Bam files were parsed into the *freebayes* v1.3.6 [75] as input for variant calling with no population prior, using two best SNP alleles, minimum mapping quality of 20, minimum coverage of 5 and theta 0.02. The VCF files were normalized based on the reference genome in *vt* v0.57721 [76]. Eigenvector and eigenvalues were created using *Plink* v2.00a3 [77] after SNPs were pruned for linkage disequilibrium (--indep-pairwise 50 10 0.1). We initially identified 7,252,950 single nucleotide variants for the whole genome and 1,773,239 single nucleotide variants within the TE regions. After the post-processing, 6,133,447 high quality SNPs were recovered from the whole genome, from which 6,130,880 were variable with 6,128,313 (99.9%) biallelic and 2,567 (0.1%) multiallelic sites. And from the TE region, 1,342,275 high quality SNPs were recovered from which 1,340,930 were variable with 1,339,585 (99.9%) biallelic and 1,345 (0.1%) multiallelic sites.

### TE content

The landscapes of TEs using Kimura 2-parameter distance were plotted using perl scripts integrated in *RepeatMasker* v4.1.0 using the chromosome assemblies. To explore TE abundance differences between genomes, Chi-square test (χ2), using the chisq.test() function in R, was performed on the number of insertions in each of the genomes and the Pearson’s residuals were plotted.

Total TE density was calculated between heterochromatin and euchromatin by intersecting masked TEs with the identified eu/heterochromatin coordinates (*bedtools* v.2.30.0 [78]). We merged masked TEs that overlapped in the Repeatmasker’s output file if they belonged to the same order, otherwise they were named ‘overlap’ using a custom script.

To calculate TE density across different genomic regions, we used the annotated gff file to extract introns, exons, upstream, downstream and intergenic regions only in the euchromatic area. Overlaps between regions were removed using *bedtools* subtract. We calculated the percentage of total length for each of the four TE orders (LTR, LINE, TIR, RC) per genomic region and in euchromatin and heterochromatin, the results were plotted using the *ggplot2* package [79]. To test if TEs are randomly distributed regarding the location to nearby genes, we performed Chi-square test statistics in R using the chisq.test() function for the total number of insertions over the size of each compartment.

To check if the sequencing and assembly metrics affected the TE content identified in each genome, we analyzed the correlation between these metrics and TE content estimates using a linear regression model.

### Identification of potentially active TEs

To find TEs that are potentially active we filtered the annotation file for TEs that have at least two full length copies in the genome. We kept those TEs that had equal to or above 99% of similarity match spanning at least 50% of the total TE consensus length using a custom bash script. We narrowed down the results for TEs that are present in at least four out of the five genomes. The final candidates were manually checked for protein coding domains, terminal repeats, and target site duplication (TSD) when applicable, following the method described in Vargas-Chavez et al. [80]. To calculate the population frequency of those active TE insertions, we used *PoPoolationTE2* v1.10.03 [81] and *TEMP2* v0.1.4 pipelines [82] since different pipelines might identify different TE insertions. The TE insertions that were detected by both *PoPoolationTE2* and *TEMP2* pipelines were merged using *bedtools merge* (*bedtools* v.2.30.0) allowing 25 bp distance (-d 25) for each insertion position.

### Cold tolerant genes

We selected a total of 26 cold tolerant associated genes using previous studies related to *D. montana* cold tolerance and the published *D. montana* genome assembly based on short reads [37–43]. We included the genes into our cold tolerant candidate gene list if they were discovered in at least three different studies (Supplementary Table S7). To transfer the coordinates to our genome, we performed a blast search of the candidate genes in the Seward chromosome-level assembly. Finally, the new coordinates of the genes were checked for homologies in *D. melanogaster* genome using *Flybase* version FB2023_01 [83].

Each gene was then scanned up to 1.5 kb up and downstream to find TEs. We focused on TEs that were found to be present in all five genomes analyzed. To check whether the selected TEs could be adding regulatory sequences, we searched for transcription factor binding sites (TFBS) and promoter motifs in TEs up-or downstream of the cold tolerant genes. We downloaded TFBS motifs related to stress response in *Drosophila* including immune response, heat shock [84,85], diapause [86–88] and cold shock domain factors [89], from JASPAR database [90] (Supplementary Table S8). We used data from *D. melanogaster* when available and otherwise from *Homo sapiens*. TFBS were identified using the web version of FIMO [91] from the MEME SUITE [92,93] with default parameters. The ElemeNT online tool was used to identify promoter motifs [94,95]

### Population analysis

A hundred and one wild-caught, individually sequenced females were used to estimate population frequencies of active TEs and TEs located nearby cold tolerant genes (Supplementary Table S3). A total of 13 populations with a minimum number of 3 individuals per population were analyzed using *PoPoolationTE2* v1.10.03 [81] and *TEMP2* v0.1.4[82]. Raw paired-end reads were trimmed using *fastp* v0.21.0 with minimum quality *Phred* score ≥ 20 (-q 20) and default parameters.

#### PoPoolationTE2

PoPoolationTE2 allows the detection of reference and de novo TE insertions in genomes. Following the manual, we first created a “TE-merged-reference” for *D. montana*, which consists of the respective masked reference genome (monSE13F37) and the TE consensus sequences generated in this work. Next, we created the “TE hierarchy file” by using an ad hoc bash script. Trimmed raw reads of each sample were mapped to the corresponding TE-merged-reference by using the local alignment algorithm *BWA* bwasw v0.7.17-r1188 [96]. Both end reads were mapped separately to the TE-merged-reference, and the paired end information was restored subsequently with module se2pe of *PoPoolationTE2*. A ppileup was generated for each sample with the *PoPoolationTE2* ppileup function (--map-qual 15). Finally, TE insertions were identified with functions identifySignatures (-- min-count 2), the final set of insertion per sample was identified with function pairupSignatures and outputs were intersected with TE coordinates located nearby cold tolerant genes using *bedtools* intersect.

#### TEMP2

To detect de novo TE insertions, we used the module insertion of *TEMP2*. We took the *RepeatMasker* annotation that was run for *PoPoolationTE2* and transformed it to bed format using gff2bed implemented in *BEDOPS* v.2.4.39 [97]. Following the *TEMP2* manual, we used *BWA mem* with options -Y and -T20 to map the paired-end reads to the corresponding reference genome. To get the fragment length of the paired-end reads, we used *Picard*’s CollectInsertSizeMetrics module v.2.26.11 [98] and inserted the mean insert size of each sample. Then, we used *TEMP2* insertion module with parameter -m 5 (percentage of mismatch allowed when mapping to TEs) to detect TE insertions.

To detect reference insertions, we used the absence module which annotates the absence of reference TE copies in the samples (parameter -x 30, the minimum score difference between the best hit and the second best hit for considering a read as uniquely mapped). We considered insertions that are in the reference genome and not annotated by the absence module as present in our samples. We identified these insertions using *bedtools* intersect with -v option. Finally, we joined the reference insertions with the ones detected by the insertion module. Outputs were intersected with TE coordinates located nearby cold tolerant genes using *bedtools* intersect.

#### Combining PoPoolationTE2 and TEMP2 TE insertions

To find a reliable set of TEs, we combined the TE insertions detected by both software. We used *bedtools* with options merge -i and -d 25 to collapse insertions overlapping or allowing a maximum distance of 25 bp between the two coordinates into a single call. We combined insertions that belonged to the same TE family.

### Chromosomal inversions

Breakpoints of one fixed and two polymorphic *D. montana* inversions were obtained from Poikela et al. [46] and Poikela et al. (in prep.) (Supplementary Table S9). Briefly, both long-and short-reads were used to identify the inversion breakpoints. PacBio and Illumina reads were mapped against Seward genome assembly (monSE13F37) using *ngmlr* v0.2.7 [99] and *BWA mem*, respectively, and the resulting bam files were parsed into *Sniffles* v2.0.7 [99] and *delly* v1.1.6 [100] structural variant identification programs, respectively. *SURVIVOR* v1.0.6 [101] was used to identify inversions that were shared by both structural variant identification programs. Finally, the putative breakpoints of all inversions were confirmed visually with the IGV using both long-and short-read data.

To check whether TEs were present in the inversion breakpoint, 50 kb regions flanking each side of the breakpoints were mapped against all five scaffold assemblies using *minimap2* to identify the breakpoint coordinates in each assembly. To validate the presence/absence of each breakpoint, we identified long reads spanning the breakpoints (3 kb on each side of the breakpoint for the two larger inversions and 500 bp on each side of the shortest inversion) using IGV. The proximal and distal positions are based on their distance to the centromere [46].

## Results

### Three new genome assemblies from *D. montana* eco-geographical distinct populations

Three new *D. montana* reference genomes from three climatically and geographically divergent populations, Crested Butte in North America (NA), Oulanka in Northern Europe (NE) and Kamchatka in Far East Asia (FE), were generated in this study (Figure 1, Table 1, and Supplementary Table S2). We also used the two other available genome assemblies, Seward and Jackson from North America *D. montana* populations [46]. As sequencing technology greatly impacts TE detection in the genome, we took advantage of long-read sequencing technique to generate these new assemblies and thus retrieve higher numbers of TEs compared to those identified based only on short-reads [102]. All five genomes were sequenced using both PacBio and Illumina technology and assembled using the same method. In North America, the populations of Jackson (monJX13F48) and Crested Butte (mon34CC5) are adapted to relatively high latitudes and high elevations (1857 m and 2960 m respectively) in the mountainous region, whereas Seward (monSE13F37) is a high-latitude coastal population (35 m). In North Europe, Oulanka population (monOU13F149) is located in northern Finland, above the arctic circle, and is adapted to cold and dark winters and short summers. The Asian sample (monKR1309) is from Kamchatka Peninsula which is a mountainous volcanic region with long cold winters in Russia (Figure 1).

**Table 1.**
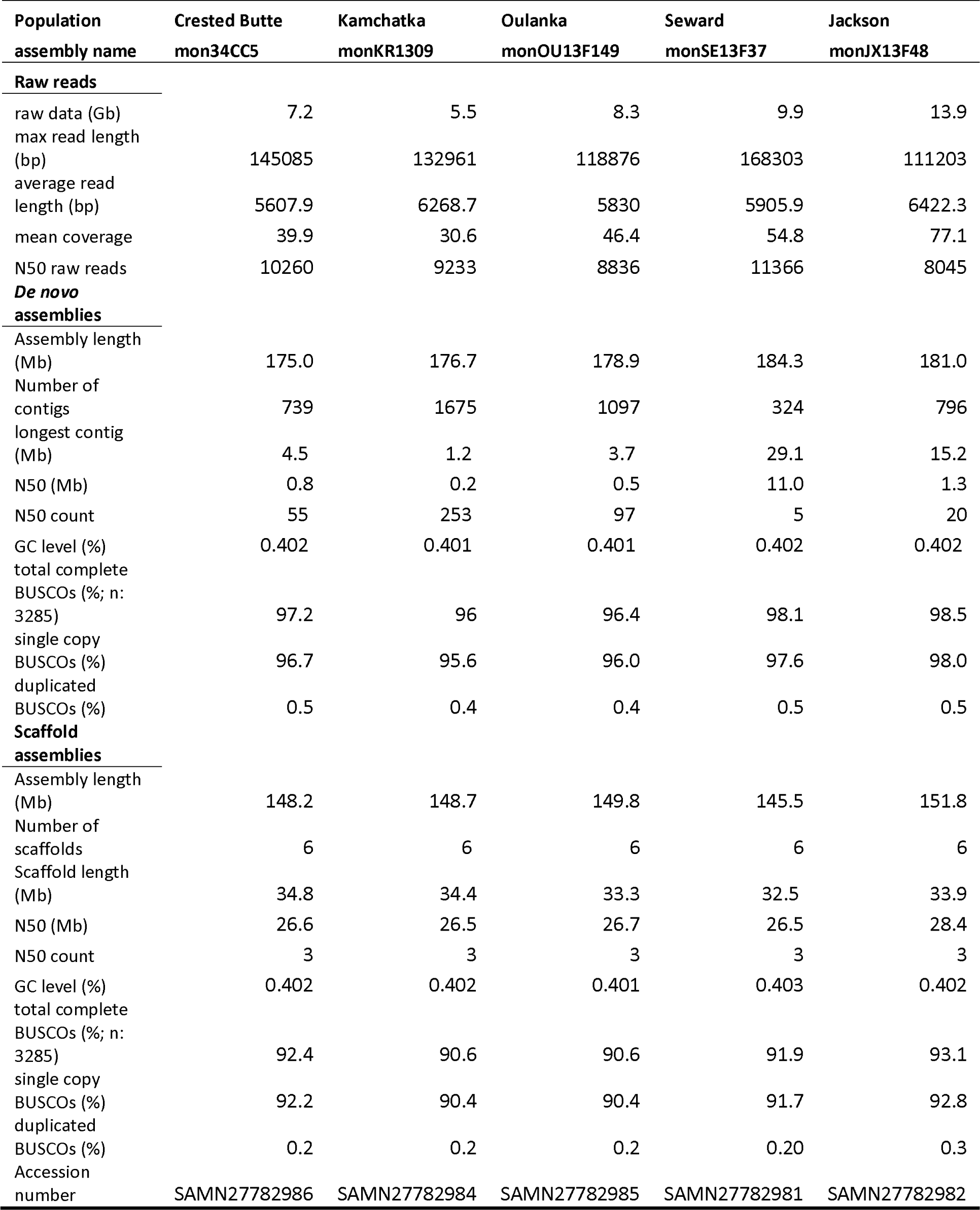
Raw read and assembly statistics for five *D. montana* genomes generated by PacBio sequencing. N50 raw reads = the read length at which half of the bases are in reads longer than or equal to this value. Raw reads metrics for Seward and Jackson are taken from Poikela et al [46].

| Population<br>assembly name | Crested Butte<br>mon34CC5 | Kamchatka<br>monKR1309 | Oulanka<br>monOU13F149 | Seward<br>monSE13F37 | Jackson<br>monJX13F48 |
| --- | --- | --- | --- | --- | --- |
| **Raw reads** |  |  |  |  |  |
| raw data (Gb) | 7.2 | 5.5 | 8.3 | 9.9 | 13.9 |
| max read length (bp) | 145085 | 132961 | 118876 | 168303 | 111203 |
| average read length (bp) | 5607.9 | 6268.7 | 5830 | 5905.9 | 6422.3 |
| mean coverage | 39.9 | 30.6 | 46.4 | 54.8 | 77.1 |
| N50 raw reads | 10260 | 9233 | 8836 | 11366 | 8045 |
| **De novo assemblies** |  |  |  |  |  |
| Assembly length (Mb) | 175.0 | 176.7 | 178.9 | 184.3 | 181.0 |
| Number of contigs | 739 | 1675 | 1097 | 324 | 796 |
| longest contig (Mb) | 4.5 | 1.2 | 3.7 | 29.1 | 15.2 |
| N50 (Mb) | 0.8 | 0.2 | 0.5 | 11.0 | 1.3 |
| N50 count | 55 | 253 | 97 | 5 | 20 |
| GC level (%) | 0.402 | 0.401 | 0.401 | 0.402 | 0.402 |
| total complete BUSCOs (%; n: 3285) | 97.2 | 96 | 96.4 | 98.1 | 98.5 |
| single copy BUSCOs (%) | 96.7 | 95.6 | 96.0 | 97.6 | 98.0 |
| duplicated BUSCOs (%) | 0.5 | 0.4 | 0.4 | 0.5 | 0.5 |
| **Scaffold assemblies** |  |  |  |  |  |
| Assembly length (Mb) | 148.2 | 148.7 | 149.8 | 145.5 | 151.8 |
| Number of scaffolds | 6 | 6 | 6 | 6 | 6 |
| Scaffold length (Mb) | 34.8 | 34.4 | 33.3 | 32.5 | 33.9 |
| N50 (Mb) | 26.6 | 26.5 | 26.7 | 26.5 | 28.4 |
| N50 count | 3 | 3 | 3 | 3 | 3 |
| GC level (%) | 0.402 | 0.402 | 0.401 | 0.403 | 0.402 |
| total complete BUSCOs (%; n: 3285) | 92.4 | 90.6 | 90.6 | 91.9 | 93.1 |
| single copy BUSCOs (%) | 92.2 | 90.4 | 90.4 | 91.7 | 92.8 |
| duplicated BUSCOs (%) | 0.2 | 0.2 | 0.2 | 0.20 | 0.3 |
| Accession number | SAMN27782986 | SAMN27782984 | SAMN27782985 | SAMN27782981 | SAMN27782982 |

Upon *de novo* assembly, the genome sizes for *D. montana* ranged from 175 to 184 Mb. By using a hybrid assembly strategy, we identified more than 95% complete single copy BUSCOs, with N50 values varying between 0.2-11.0 Mb (Table 1). After scaffolding, the total genome size and BUSCO values decreased (90.6 – 93.1%, 145-151 Mb) compared to the non-scaffolded assemblies. This is because we were unable to assign all contigs to chromosome scaffolds and the unassigned contigs were excluded from the final chromosome-level *D. montana* genome [46]. The detailed information on raw reads, contigs and scaffolds statistics are given in Table 1. We used complete genome assembly for TE identification and chromosome level assemblies excluding the heterochromatic regions (see below) for the rest of our analyses in this work.

### Over ninety percent of the TE consensus sequences identified in *D. montana* belong to new families or new subfamilies

By using genomes from climatically and geographically diverged *D. montana* populations, we wanted to include as much intraspecific diversity into our comprehensive TE library as possible. We used *REPET* to *de novo* annotate TEs in each of the five genomes. On average, 3,221 TE consensus were built per genome, and 303 consensus per genome were kept after manual curation (Supplementary Table S10). After clustering individual libraries, a total of 555 non-redundant TE consensus sequences were recovered (TE library is provided as Additional file 4). Ninety two percent of the TE consensuses are described as new families (445) and new subfamilies (66) and only the remaining 8% corresponds to previously known families (44) (Figure 2 and Supplementary Table S11). Long terminal interspersed repeats (LTR) are the most abundant TE order in the *D. montana* library of which, Gypsy superfamily is the most abundant (80%) followed by Bel-Pao (14%). LINE and DNA transposons (TIR, terminal inverted repeat, and RC, rolling circle elements) contribute almost equally to the TE library (12%) with the highest abundance of Jockey (68%) and Tc1-Mariner (40%) in each order, respectively (Figure 2). We used the manual curated library to annotate each of the five reference genomes and analyzed the abundance, distribution, and activity of TE sequences. We first analyzed whether differences in sequencing or assembly metrics explained variation in the TE metrics estimated (Supplementary Table S12). The N50 of contigs had a significant effect on the total number of raw TE sequences discovered in the heterochromatin, while scaffold statistics did not impact any of the TE metrics analyzed (Supplementary Table S12). Because all of the following analyses have been made on the euchromatin, our results are thus not affected by sequencing or assembly differences across genomes.

**Figure 2.**
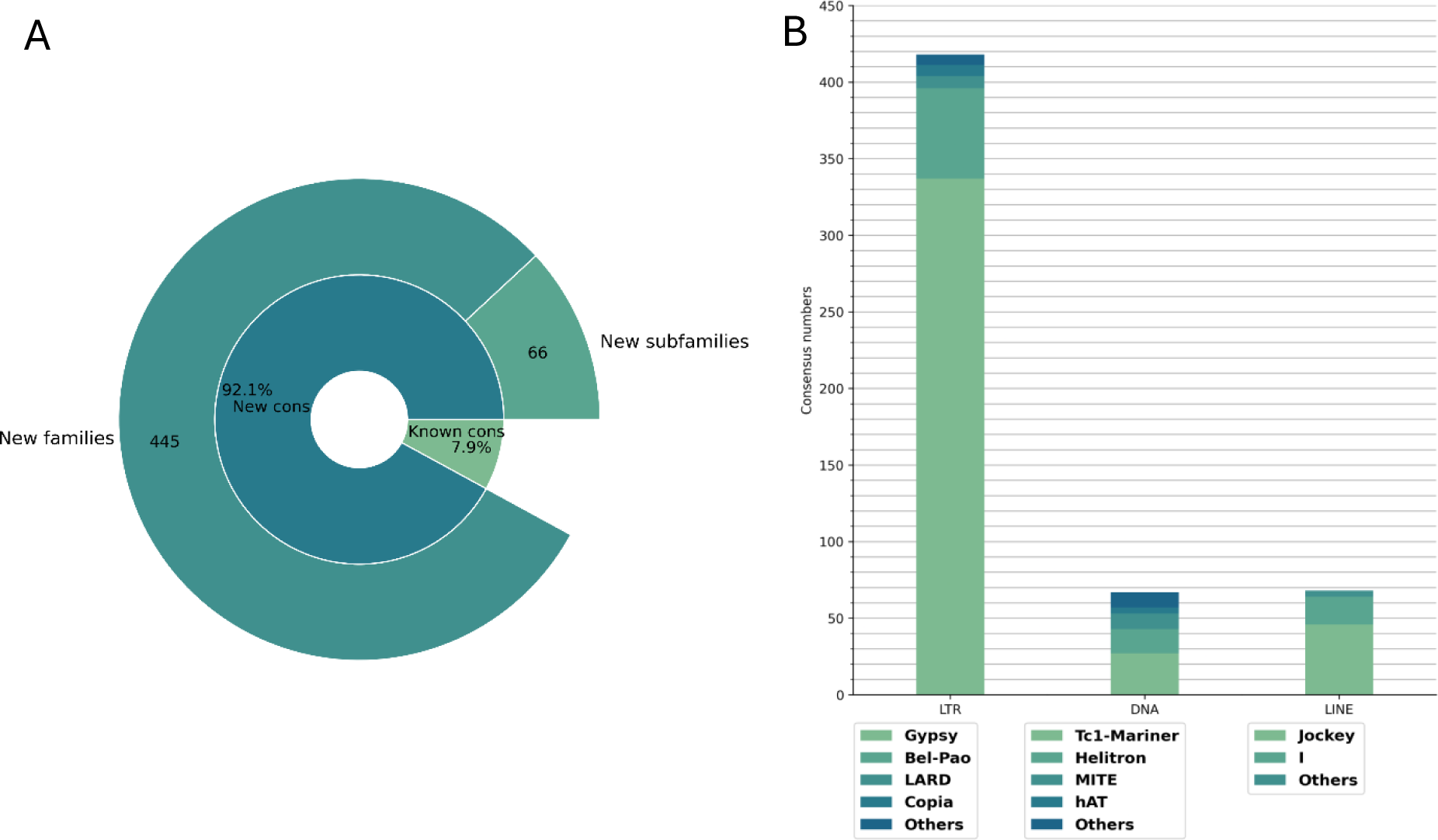
Consensus sequence classifications of the curated TE library built from *D. montana* genomes. **(A)** Percentage of new to known TE consensus (cons) sequences. **(B)** Numbers of consensus elements classified as LTR, DNA (RC and TIR) and LINEs.

We also explored the genetic variation across the whole genome and within TE sequences in the five *D. montana* samples by performing a principal component analysis (PCA) with biallelic SNPs. Results from the two PCA analyses are highly concordant, with the first PC separating the three NA populations from the Oulanka (NE) and Kamchatka (FE) populations and explaining similar amounts of variation: 33 and 32% for the whole genome and TE region SNPs, respectively (Figure 3A & 3B and Supplementary Table S13). The second PC, which explains 28 and 29% of the variation, separates the Crested Butte population (NA) from the other two NA populations: Jackson and Seward. This pattern likely reflects the demography of these populations (Poikela et al., in prep) and shows the high level of intraspecific variation in the *D. montana* genomes analyzed.

**Figure 3.**
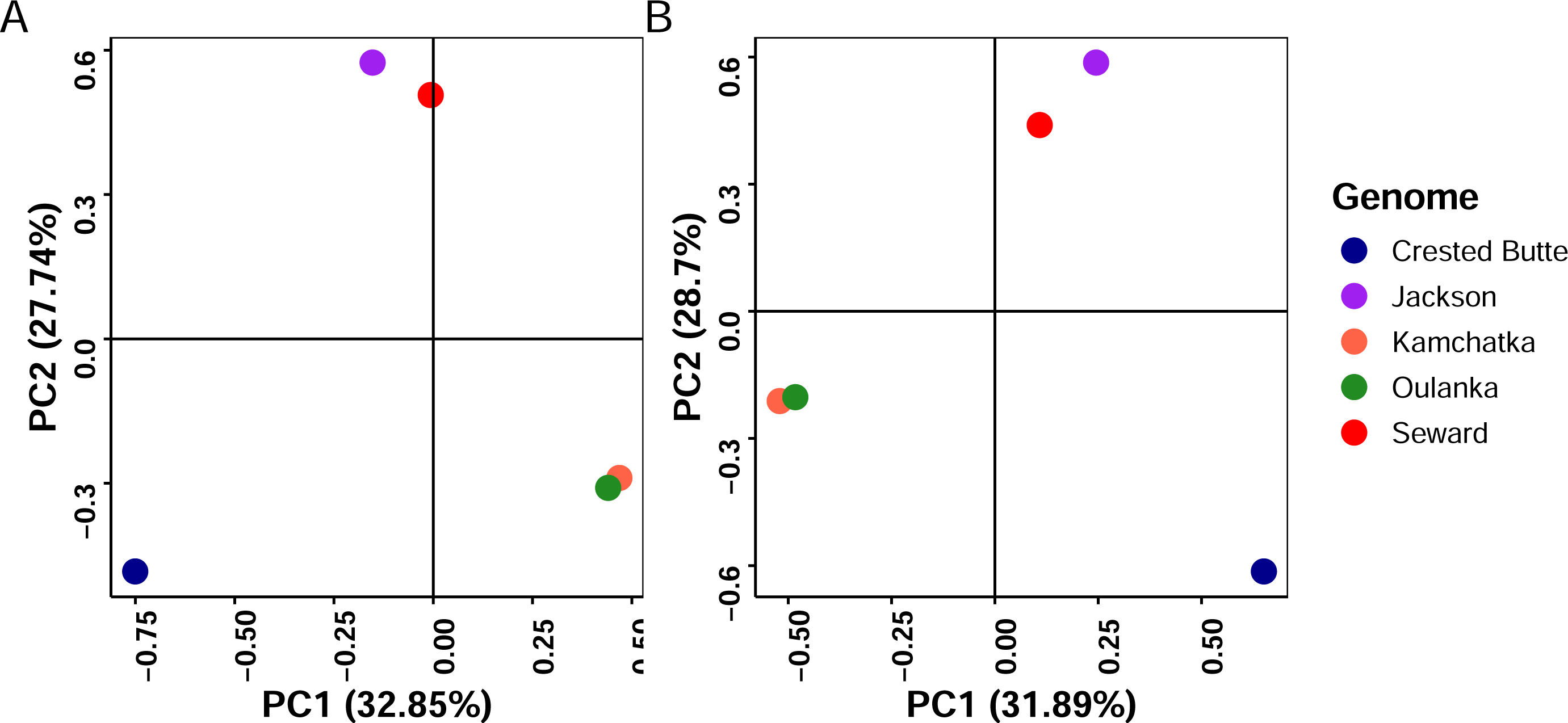
Principal component analysis (PCA) on SNPs **(A)** across the whole genome and **(B)** inside TE regions in five *D. montana* samples originating from different populations.

### Transposable elements contribute 11% to 13% to the *D. montana* genomes

Whole genome TE content across genomes varied between 10.7% and 13% (Table 2). In all five genomes, LTRs were the most abundant order, followed by rolling circles (RC), LINEs, and terminal inverted repeats (TIRs) (Figure 4A). TE abundance at the order and superfamily level varied across genomes (Figure 4B and 4C). In Seward, LTR (χ^2^ residue -8.2, p-value < 0.05) is the least abundant and RC (χ^2^ residue 5.7, p-value < 0.05) is the most abundant order compared with the other genomes analyzed. At the superfamily level, Seward shows the highest variation in TE content; Gypsy and Bel-Pao have less insertions compared with other genomes (χ^2^ residue -11.1 and -6.9, p-value < 0.05), while Helitron, Maverick (RC) and LARD have more insertions compared with other genomes (χ^2^ residue 4.4, 4.2, and 5.7, p-value < 0.05). Gypsy has the highest number of insertions in Crested Butte (χ^2^ residue 5.2 p-value < 0.05), followed by Kamchatka (χ^2^ residue 3.2 p-value < 0.05), while LARD has the lowest insertion number in Crested Butte compared to other genomes (χ^2^ residue -6.8, p-value < 0.05) (Figure 4C). Overall, while RC elements did not show a lot of variability at the library level (Figure 2B) they annotate a considerable number of copies in the five genomes (Figure 4A).

**Figure 4.**
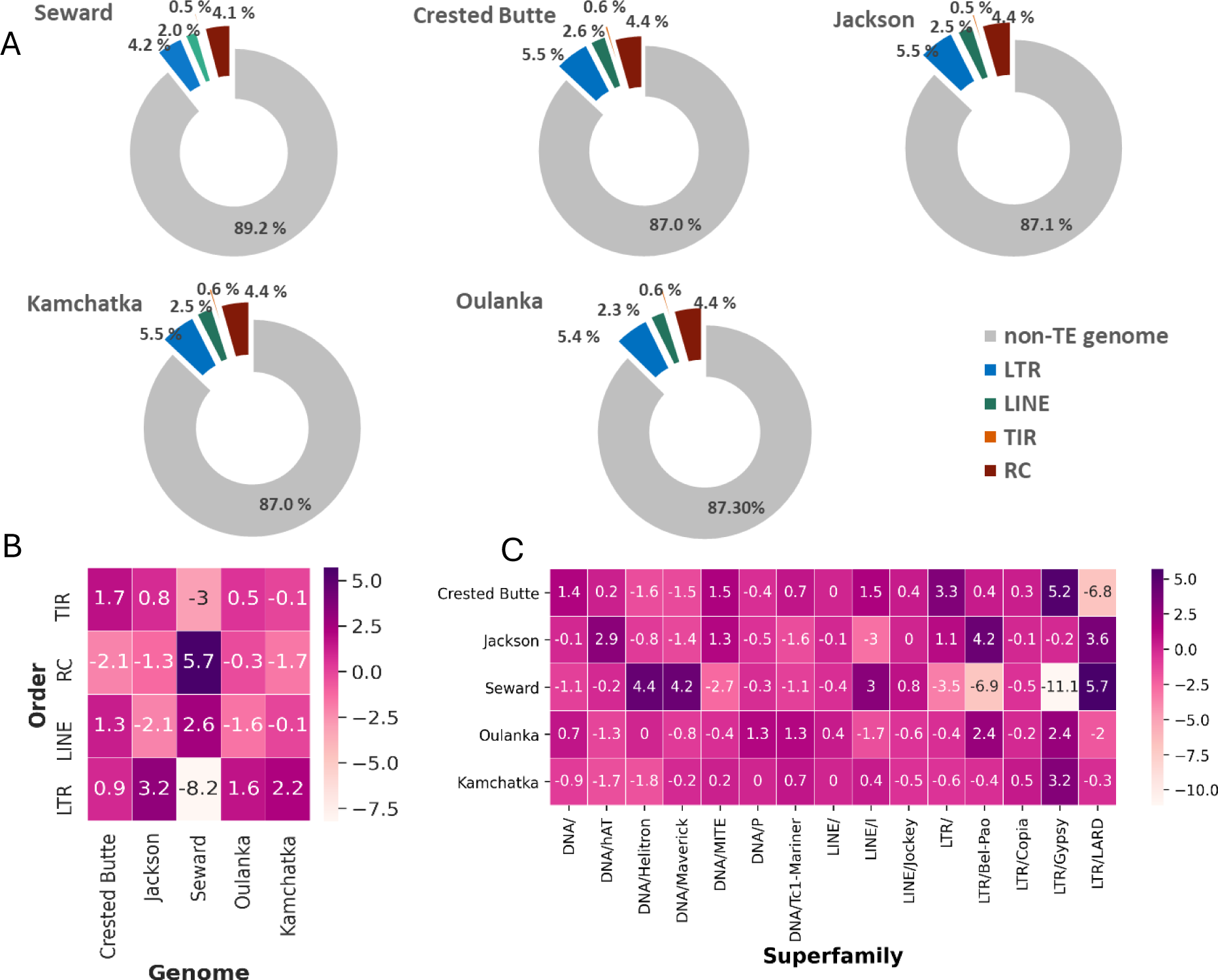
TE content in the five *D. montana* genomes analyzed. **(A)** Percentage of TE orders occupying each genome. **(B)** Heatmap showing the Pearson residuals in different populations based on TE copy numbers at the order, and **(C)** the superfamily levels. Superfamilies with less than 50 copies are grouped into DNA/ LTR/ and LINE/.

**Table 2.**
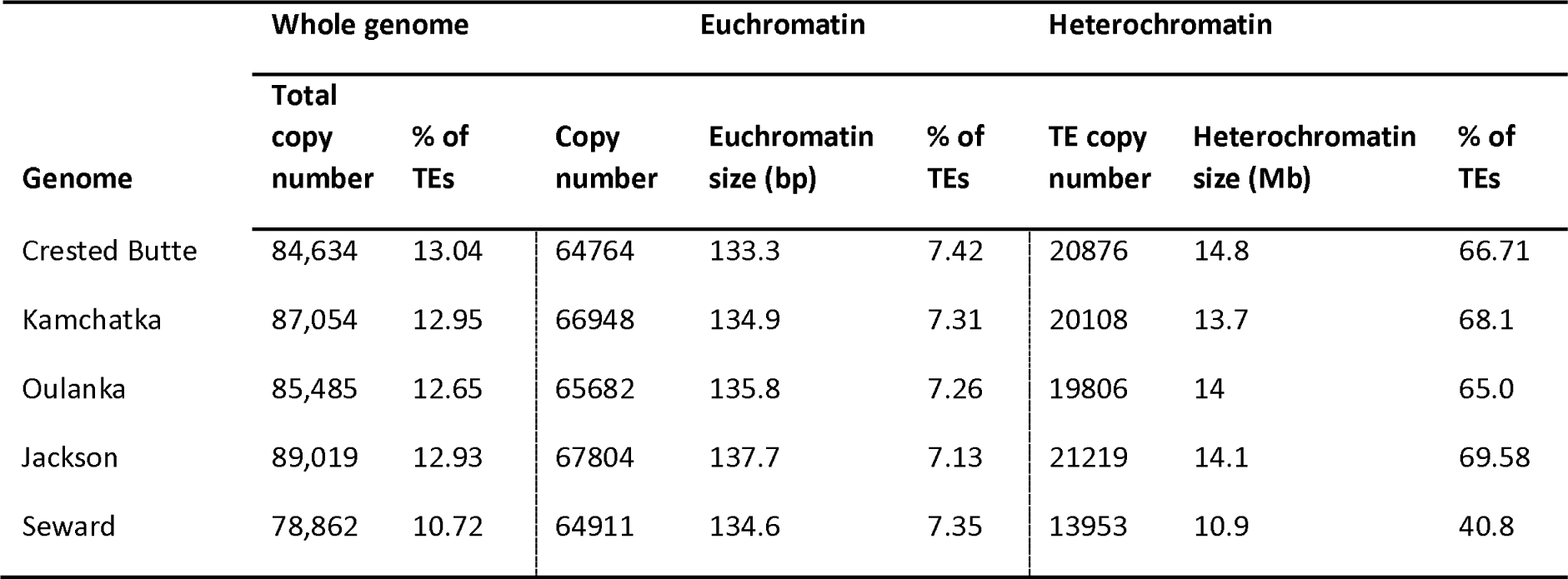
Description of TE copy numbers annotated in the whole genomes and in eu-and heterochromatin using RepeatMasker v4.1.0.

| Genome | Whole genome |  | Euchromatin |  |  | Heterochromatin |  |  |
| --- | --- | --- | --- | --- | --- | --- | --- | --- |
|  | Total copy number | % of TEs | Copy number | Euchromatin size (bp) | % of TEs | TE copy number | Heterochromatin size (Mb) | % of TEs |
| Crested Butte | 84,634 | 13.04 | 64764 | 133.3 | 7.42 | 20876 | 14.8 | 66.71 |
| Kamchatka | 87,054 | 12.95 | 66948 | 134.9 | 7.31 | 20108 | 13.7 | 68.1 |
| Oulanka | 85,485 | 12.65 | 65682 | 135.8 | 7.26 | 19806 | 14 | 65.0 |
| Jackson | 89,019 | 12.93 | 67804 | 137.7 | 7.13 | 21219 | 14.1 | 69.58 |
| Seward | 78,862 | 10.72 | 64911 | 134.6 | 7.35 | 13953 | 10.9 | 40.8 |

### Transposable elements are not randomly distributed across the genome

As expected, the percentage of TEs in euchromatin (∼7%) was much lower than the percentage of TEs in heterochromatin (41%–68%; χ^2^ test p-value < 2.2×10, Supplementary Table S14A, Figure 5A and Table 2). At the chromosomal level, the highest proportion of TEs were found in chromosome 4 and chromosome X while chromosome 2R has the lowest proportion (Figure 5B), which is correlated with the chromosome size (Spearman Correlation Rho = 0.9123, p-value = 1.711×10). However, when comparing the TE content in autosomes versus the X chromosome, the X chromosome had higher TE density than autosomes (Wilcoxon test p-value = 1.403x10; Figure 5C). Finally, the distribution of TEs regarding the nearest gene position showed that TEs are more abundant in intergenic regions, while exons bear the lowest percentage (Figure 5D and Supplementary Table S14B) (χ^2^ test p-value < 2.2×10) as expected because the majority of TE insertions in exons are likely to be deleterious.

**Figure 5.**
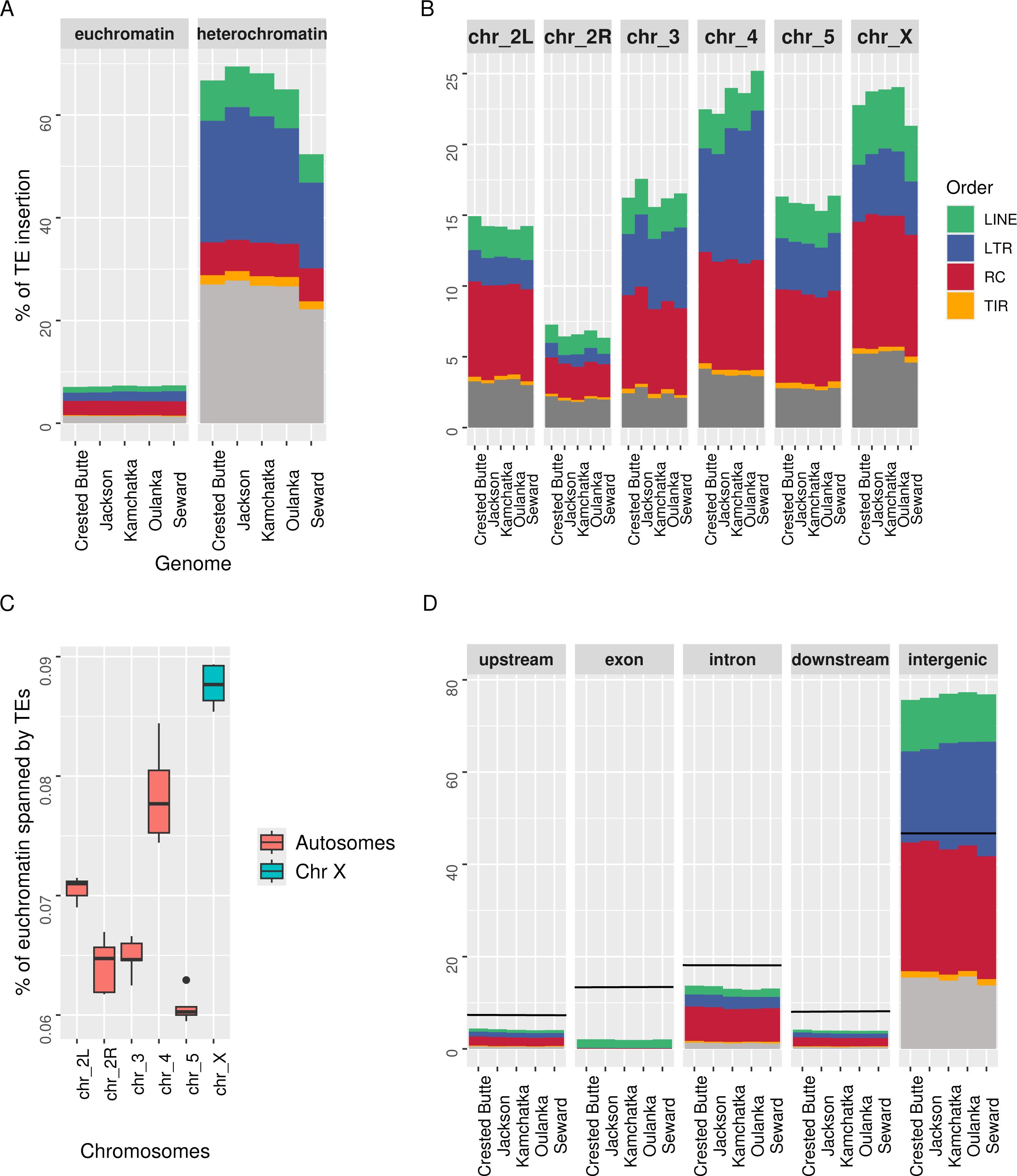
TE distribution in five *Drosophila montana* genomes analyzed. **(A)** Percentage of eu-and heterochromatin regions occupied by TEs, normalized by the total region’s length. **(B)** Percentage of TEs per chromosome, normalized by the total chromosome length. **(C)** Boxplot showing the percentage of TEs in euchromatin in each chromosome. Autosomes are shown in turquoise, and the X chromosome in red. **(D)** Percentage of TEs (of different orders) per genomic region, normalized by the length of each region. The black lines indicate the expected percentage. ‘Overlap’ means that TEs having overlapping coordinates but do not belong to the same order.

### Most of the potentially active TEs in *D. montana* are retrotransposons

We compared the TE landscapes, *i.e.* the proportion of each TE superfamily plotted against the genetic distance observed between the TE copies and the consensus sequences, for the five genomes analyzed. The TE landscapes showed periods of transposition bursts in all five genomes and depicted a bimodal curve (Figure 6A) [103]. The first peak showed the highest enrichment of young elements, mostly LTR/Gypsy (Figure 6A). The second peak, between Kimura distance 10-15, is a burst of LTR/LARD families. Helitrons dynamics follow a bell curve consistent with the equilibrium mode of transposition-excision over a very long period of time [103].

**Figure 6.**
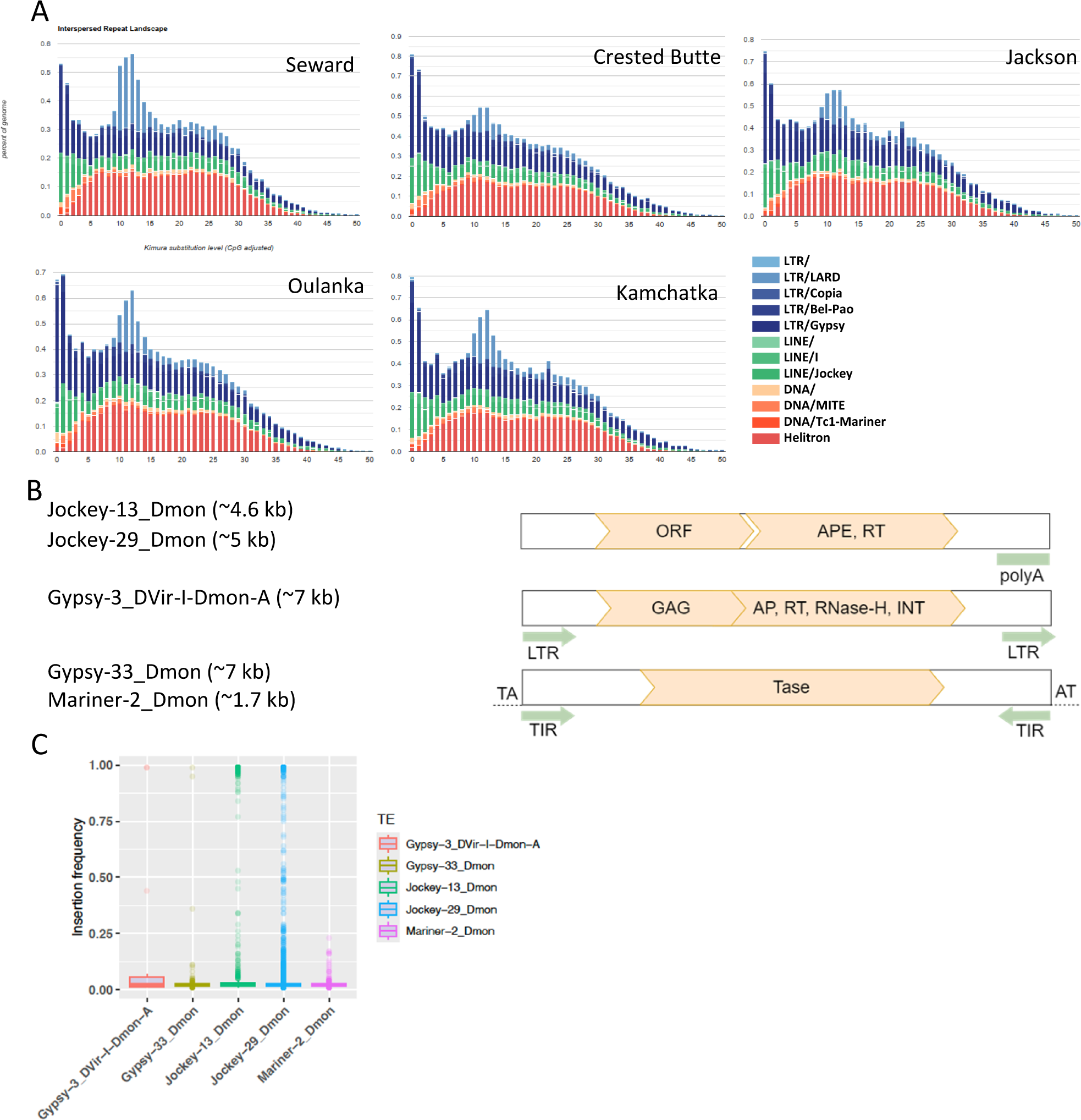
Identification of potentially active TEs in the five *D. montana* genomes analyzed. **(A)** Landscape of TEs in the five genomes **(B)** Schematic representation of the structure of potentially active TE families identified in *D. montana*. APE: apurinic endonuclease, RT: reverse transcriptase, INT: integrase, AP: aspartate proteinase, Tase: transposase, LTR: long terminal repeats, TIR: terminal inverted repeats. **(C)** Boxplot showing insertion frequencies detected by both PoPoolationTE2 and TEMP2 softwares from 13 population data sets, the upper dots are outliers.

We further investigated whether any of the TE families identified could be potentially active. We found five potentially active TE families in *D. montana* including two Jockey, two Gypsy and one Tc1-Mariner superfamilies (Figure 6B). Copies from these active elements were found in the majority of genomes: *Jockey-29_Dmon* and *Mariner-2_Dmon* were not present in the Kamchatka genome, *Gypsy-3_DVir-I-Dmon-A* and *Jockey-13_Dmon* were not present in Oulanka, and *Gypsy-33_Dmon* was not present in Seward. To estimate the frequency of the active TEs in euchromatic regions, we checked the frequency of each insertion across 13 populations of *D. montana* using *PoPoolationTE2* and *TEMP2* pipelines (Figure 1 & 6C). We considered only those insertions that were detected by both PoPoolationTE2 and TEMP2 pipelines (see Material and Methods). The most abundant potentially active families were *Jockey-29_Dmon* (1,925 copies), followed by *Jockey-13_Dmon* (779 copies). The number of copies found for *Mariner-2_Dmon*, *Gypsy-33_Dmon*, and *Gypsy-3_Dvir-I-Dmon-A* were 166, 134, and 14, respectively. The population frequencies of the five active TEs are predominantly below 0.1, consistent with these insertions being recent (Supplementary Table S15). Two of the families, *Jockey-13_Dmon* and *Jockey-29_Dmon* had copies present at high frequencies in the genome, suggesting that some of these copies could have increased in frequency due to positive selection.

### TE insertions could affect the regulation of cold tolerant genes in *D. montana*

To investigate if TEs may have an adaptive role in the species adaptation to the northern climate, we searched for TEs located in or nearby the 26 previously identified cold tolerant candidate genes (Supplementary Table S7). We found a total of 649 TE insertions belonging to 44 families that were located inside or nearby candidate genes. Out of 649 TE insertions, 34 TEs were found in all five genomes analyzed located inside or nearby 13 cold-tolerant genes (Table S16). Additionally, we found 66 TEs present in 2-5 genomes and 6 TEs present in only one of the genomes analyzed, suggesting that these TE copies could be important only in some or in one of the populations, which could be related to the eco-geographical differences across populations (Supplementary Tables S17 & S18).

To investigate if the 34 TEs present in all five genomes and located nearby 13 out of 26 cold tolerant genes might have regulatory effects on these genes, we searched for promoter elements and transcription factor binding site (TFBS) motifs in TEs located up-and downstream of each gene and for TFBS motifs in TEs located in the first intron. We searched for TFBS relevant for cold adaptation, *i.e.* stress response (cold, immune and heat shock) and diapause (see Material and Methods). We found that one Gypsy, one LARD, and one Helitron family have possible active promoter elements upstream of *Mlc2*, *Inos* and *CG12057* genes, and two Helitrons have possible active promoter elements downstream of *Yp2* and *Ltn1* genes in all five genomes (Figure 7 and Supplementary Table S19). In the Kamchatka population, a Helitron insertion downstream of gene *CG17571* also contains promoter motifs (Supplementary Table S19). Additionally, we found TFBS motifs related to immunity response and diapause (q < 0.5) in TEs located nearby six cold tolerant genes (Supplementary Table S20). A Diapause related FOXO motif; FOXO1::ELF1 [104], had the highest quality match (q < 0.05) found in one R1 and one Helitron element downstream of *blow* and *Yp2* genes, respectively (Figure 7, Supplementary Table S20). More FOXO motifs were detected in TEs upstream of genes *CG12057* and *CG17571*. Moreover, the immunity response factor, nub [105] was detected in the Helitron and Gypsy elements located in the first intron of *Abl* and *CG12576*, respectively, and in the Helitron upstream of *CG12057* in three of the five populations (Supplementary Table S20). In *Abl*, one stress-related motif, su(Hw) [106] and two motifs related to diapause (Abd-B, ems) were found in a series of elements located in the first intron (Figure 7, Supplementary Table S20).

**Figure 7.**
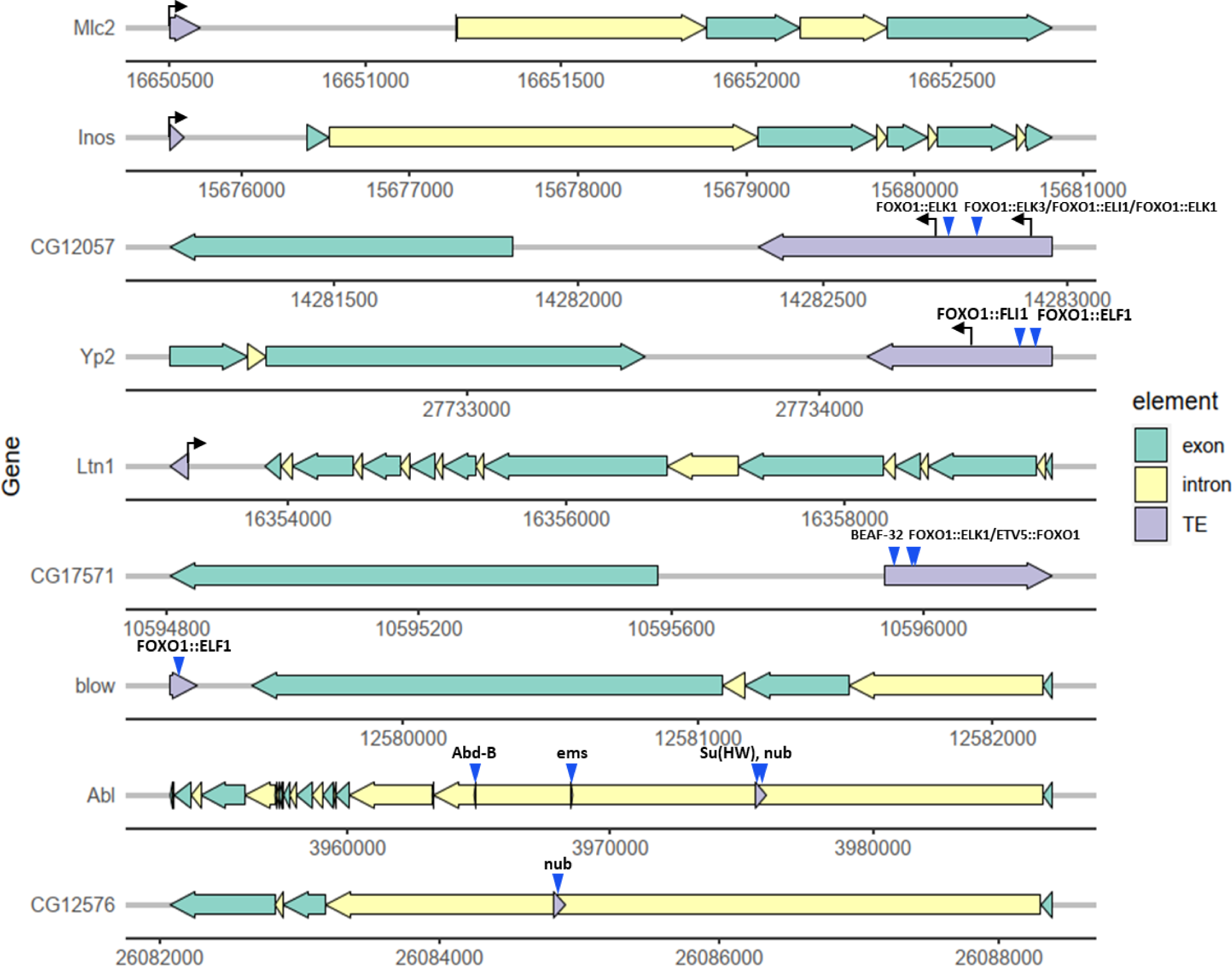
TEs present in the five *D. montana* genomes located upstream, downstream, or in the first intron of cold-tolerant genes. Arrows represent the promoters and triangles the TFBS.

Finally, we estimated the frequency of the 34 TEs present in the five genomes in 13 populations from Europe and America (Figure 1). If these TEs are likely to be adaptive, we would expect them to be present at high frequencies in at least some of the populations analyzed. *PopoolationTE2* failed to detect eight of the TE insertions. On the other hand, *TEMP2* identified all 34 TE insertions across populations, with all of them except two insertions being fixed in the 13 populations analyzed (Supplementary Table S21).

Overall, we found TEs containing regulatory sequences and present at high population frequencies or fixed, and thus more likely to have a functional role compared with TEs present at low population frequencies, located nearby nine of the 26 previously identified candidate genes involved in cold adaptation in *D. montana* (Figure 7).

### TEs are present in the breakpoints of the three *D. montana* inversions analyzed

We searched for TE insertions at breakpoints of three *D. montana* inversions: one in chromosome X present in the five genomes analyzed and two polymorphic inversions in chromosome 4 (Table 3 and Supplementary Figure S2). These inversions are described in Poikela et al. ([46], and in prep). The chromosome X inversion is a fixed inversion compared to *D. flavomontana*. One of the polymorphic inversions in chromosome 4 is present only in the Crested Butte genome while the other one is a short inversion of only 1.5 kb and it is found in the genomes of Oulanka and Kamchatka (Poikela et al, in prep).

**Table 3.** Transposable elements that were found in the breakpoint regions (3 kb /500 bp) of the three inversions analyzed in this work. TEs that are present only in the genomes with inversion breakpoints (and not in other genomes) are in bold.

| Chr | Inversion coordinates |  | Genomes | TE content |  |
| --- | --- | --- | --- | --- | --- |
|  | Proximal | Distal |  | Proximal breakpoint | Distal breakpoint |
| X | 11174369 -<br>11174380 | 3994112 -<br>3994127 | Seward | R1-2_Dmon |  |
|  |  |  |  | Maverick-1_Dmon |  |
|  |  |  |  | TART_DV-Dmon-B |  |
|  |  |  |  | **Helitron-5_Dmon** |  |
|  |  |  |  | **Gypsy-102_Dmon** |  |
|  |  |  |  | **Helitron-10_Dmon** |  |
|  |  |  |  | **Helitron-2N1_Dvir-Dmon-B** |  |
|  |  |  |  | **Helitron-2N1_Dvir-Dmon-A** |  |
|  |  |  |  | **Helitron-1N1_Dvir** |  |
|  | 11136142 -<br>11136155 | 3936207 -<br>3936246 | Jackson | R1-2_Dmon |  |
|  |  |  |  | **Helitron-5_Dmon** |  |
|  |  |  |  | **Gypsy-102_Dmon** |  |
|  |  |  |  | **Helitron-10_Dmon** |  |
|  |  |  |  | Helitron-5_Dmon |  |
|  |  |  |  | **Helitron-2N1_Dvir-Dmon-B** |  |
|  |  |  |  | **Helitron-2N1_Dvir-Dmon-A** |  |
|  |  |  |  | **Helitron-1N1_Dvir** |  |
|  | 11110783 -<br>11110796 | 3984908 -<br>3984929 | Kamchatka | R1-2_Dmon |  |
|  |  |  |  | Maverick-1_Dmon |  |
|  |  |  |  | **Helitron-5_Dmon** |  |
|  |  |  |  | **Gypsy-102_Dmon** |  |
|  |  |  |  | **Helitron-10_Dmon** |  |
|  |  |  |  | Helitron-5_Dmon |  |
|  |  |  |  | **Helitron-2N1_Dvir-Dmon-B** |  |
|  |  |  |  | **Helitron-2N1_Dvir-Dmon-A** |  |
|  |  |  |  | **Helitron-1N1_Dvir** |  |
|  | 11304271 -<br>11304284 | 4034972 -<br>4035011 | Oulanka | Maverick-1_Dmon | Maverick-1_Dmon |
|  |  |  |  | **Helitron-5_Dmon** | Maverick-1_Dmon |
|  |  |  |  | **Gypsy-102_Dmon** |  |
|  |  |  |  | **Helitron-10_Dmon** |  |
|  |  |  |  | Helitron-8_Dmon |  |
|  |  |  |  | **Helitron-2N1_Dvir-Dmon-B** |  |
|  |  |  |  | **Helitron-2N1_Dvir-Dmon-A** |  |
|  |  |  |  | **Helitron-1N1_Dvir** |  |
|  | 10289870 -<br>10289883 | 3549579 -<br>3549600 | Crested Butte | R1-2_Dmon | Maverick-1_Dmon |
|  |  |  |  | **Helitron-5_Dmon** |  |
|  |  |  |  | **Gypsy-102_Dmon** |  |
|  |  |  |  | **Helitron-10_Dmon** |  |
|  |  |  |  | Helitron-5_Dmon |  |
|  |  |  |  | Helitron-1N1_Dvir |  |
|  |  |  |  | **Helitron-2N1_Dvir-Dmon-B** |  |
|  |  |  |  | **Helitron-2N1_Dvir-Dmon-A** |  |
|  |  |  |  | **Helitron-1N1_Dvir** |  |
| 4 | 6596948 -<br>6599447 | 7298775 -<br>7298610 | Crested Butte | **Gypsy-5_DVir-I-Dmon-F** | **MITE-1_Dmon** |
|  |  |  |  | **Gypsy-40_Dmon** | Helitron-1_Dvir-Dmon-C |
|  |  |  |  | Gypsy-5_DVir-I-Dmon-A | **Gypsy-215_Dmon** |
|  |  |  |  | **MITE-1_Dmon** | Gypsy-181_Dmon |
| 4 | 2906226 | 2907786 -<br>2907896 | Oulanka | Helitron-4_Dmon<br>Gypsy-83_Dmon | Gypsy-70_Dmon<br>**Helitron-5_Dmon**<br>**Gypsy-8_DVir-I-Dmon-B**<br>ULYSSES_I-Dmon-H<br>Helitron-4_Dmon<br>Helitron-1_Dvir-Dmon-C |
|  | 3165124 -<br>3165142 | 3166689 | Kamchatka | Helitron-4_Dmon<br>Gypsy-83_Dmon | Gypsy-70_Dmon<br>**Helitron-5_Dmon**<br>**Gypsy-8_DVir-I-Dmon-B**<br>Helitron-4_Dmon |

For the fixed inversion on chromosome X, we found R1, Maverick, TART, Helitron, and Gypsy elements in the proximal breakpoint. From these elements, Gypsy and Helitron insertions were found in all five genomes, while R1 was found in four of them (Table 3). Only Maverick elements were found in the distal breakpoint in two of the genomes (Table 3). The longer 4th chromosome polymorphic inversion, present only Crested Butte genome, contain Gypsy and MITE elements in both the distal and the proximal breakpoints. The breakpoints of the shorter 4th chromosome polymorphic inversion, present in Oulanka and Kamchatka genomes, contained Helitron and Gypsy elements both in the proximal and the distal breakpoints (Table 3).

## DISCUSSION

*Drosophila montana* is well-adapted to northern environments, which has enabled its populations to expand in near-arctic and arctic regions where winter spans almost half of the year at sub-zero temperature, and summer is short occasionally including hot periods. Multiple studies have investigated *D. montana*’s adaptation to northern environments at the genetic level [34,37,38,40,42,107]. Although transposable elements (TEs) have been shown to play a role in environmental adaptation [18], nothing is known about the potential role of TEs in *D. montana* adaptation to cold environments. By using flies across different continents and generating *de novo* genome assemblies with long-reads (PacBio), we built the first comprehensive TE library for *D. montana* and utilized it to investigate TE dynamics in this species and the association between TEs and adaptation to harsh northern environments.

### *D. montana* genomes contain 445 new TE consensus sequences

After several steps of stringent filtering to reduce false positives and redundancy, we identified a high number of new TE consensus sequences in *D. montana*, 92% of the total TE consensus, mostly belonging to the LTR order and Gypsy superfamily (Figure 2). Only 8% of the TE consensus sequences corresponded to known TE families, mostly to *D. virilis* annotations, which is more closely related to the studied species than *D. melanogaster* (Supplementary Table S11). There are several potential reasons for the high number of new TE families. Probably the most important reason is that *D. montana* is highly diverged from the other model and non-model *Drosophila* species for which TE libraries exist in databases (between ∼40 MYA from *D. melanogaster* and 9 MYA from *D. virilis*) [108,109]. Another plausible reason comes from the species’ unique distribution in the Holarctic region. Upon colonization into new habitats, species can be exposed to the invasion of new TEs through horizontal transfer from new parasites and from other conspecies [110]. Similar events could have happened in *D. montana* while occupying new cold habitats.

Our principal component analysis (PCA) on genome-wide distribution of SNPs and within TE regions shows the high level of intraspecific variation that we aimed to include into our TE library (Figure 3). The PCA depicts the high divergence not only between *D. montana* populations from North America and Northern Europe, but also among North American populations as Crested Butte (Colorado) is highly diverged from Jackson (Wyoming) and Seward (Alaska). High divergence of Crested Butte and other North American populations is likely a consequence of past founder effects potentially associated with genetic drift, variable selection pressures, and local adaptation [43]. Moreover, the large inversion on chromosome 4 is uniquely fixed in Crested Butte (Poikela et al., in prep.), which further reduces genetic exchange between populations and results in elevated genetic divergence (Poikela et al., in prep. and [27]). Overall, environmental heterogeneity through time and space can act as a selective force driving adaptive differentiation between populations. Interestingly, Kamchatka shows high similarity to the Finnish population from Oulanka regardless of their geographic distance. This affinity could be explained by the ancestral route of expansion from NA populations towards Eurasia through the Bering Strait [35], therefore, the genetic divergence of European and Asian populations would be low.

### TE content in *D. montana*

Our *de novo* TE annotations confirmed that the amount of TE content significantly contributes to the *D. montana* genome size as reported for other species [10,111]. The overall average TE content in *D. montana*, 11-13%, is similar to that of its close relatives, *D. virilis* (15%) [11] and *D. incompta* (13-14%) [103], and smaller than those of *D. melanogaster* (20%) [111] and *D. suzukii* (30%) [11]. LTRs represent the highest genomic fraction of ∼5% followed by RC elements (Helitron and Maverick), ∼4%, LINE, ∼2%, and finally TIR with below 1% (Figure 4A). According to this observed pattern, LTRs are the most abundant elements which is typical for many *Drosophila* flies, however, the abundance does not follow the assumed global pattern: LTRs>LINEs>TIRs>OTHERs [10,11].The higher Helitron fraction in *D. montana* genome (compared to the global pattern) combined with its rather persistent activity spanned over an extended course of time (Figure 6A) suggests that Helitrons have been tolerated more than other elements probably due to their neutral or adaptive effects. The relaxed selective pressure on Helitron transposition could come from the element’s own nature and its tendency to insert near less critical genes.

In *D. montana* a significantly higher number of TEs were accumulated in the heterochromatin region. The heterochromatin’s resilience to TE accumulation is due to the scarcity of active genes, and as expected, TEs were less tolerated in the gene-rich region or the euchromatin (Figure 5A). According to the *Drosophila* 12 Genomes Consortium (2007), 1-9% of the euchromatin is occupied by TEs across *Drosophila* species, and hence, the calculated 7% for *D. montana* is within the reported range. Analysis of the TE content distribution between different genomic regions reveals higher accumulation of TEs in the intergenic area followed by introns, with the least occupancy in exons (Figure 5D). Overall, TE distribution was not random throughout the genome or between chromosomes suggesting that purifying selection acts against TE insertions due to their multiple deleterious effects [112]. TE density was significantly higher in chromosome 4 and X and the lowest in the right arm of chromosome 2 (2R). Apart from the chromosome sizes, the higher acquisition of TEs on sex chromosome could be associated with dosage compensation [113]. Moreover, Wiberg et al. [43] found that the top candidate SNPs for cold adaptation in *D. montana* are enriched on chromosomes 4 and X which also include several inversions (see below).

According to the landscape of TEs, all five genomes show similar dynamics of the TE content. We found that the *D. montana* TE landscape is comparable to that of *D. melanogaster* which also has a large fraction of young and active LTR elements, while it differs from the *D. simulans* landscape which instead has a large fraction of old and degraded elements [11]. The pattern is bimodal with a general trend towards a small number of highly diverged/old elements (Figure 6A). The dynamic pattern of TE landscapes is often a reflection of the species demographic history [114,115]. For example, occupying new environments will pose various types of stress to the species of which, cold is one stress factor in the northern regions, which can potentially re-activate mobile elements [12,116]. The bimodal pattern indicates two bursts of transposition events that have interrupted the transposition/excision equilibrium (standard bell shape pattern). This pattern is generally seen in specialist species and species with small effective population size which happens when the distribution is patchy and limited by environmental resources [103]. The five genomes also have similar numbers of potentially active TEs (Figure 6B), which are mostly present at low populations frequencies (Figure 6C) as expected if these are young insertions.

### TEs associated with cold tolerant genes

Our analysis of the TEs located nearby cold-tolerant genes in the five genomes suggest that some of them might be playing a role in the regulation of these genes as they are present at high population frequencies and contain regulatory regions (Figure 7 and Supplementary Table S21). Three types of FOXO1 and one BEAF-32 TFBS, among other TFBS, were found inside some of these TEs (Figure 7). *Helitron-2N1_Dvir-Dmon-B* located downstream of *Yp2* and upstream of *CG12057* also has promoter motifs and FOXO transcription factors. FOXO transcription factors regulate cellular homeostasis, longevity and response to stress [117]. In *D. montana*, activation of FOXO has been suggested to be connected with the flies’ overwintering ability [86]. Moreover, *Helitron-1_Dvir-Dmon-C* upstream of *CG17571* has a match to promoter elements only in the Kamchatka genome. This TE matches BEAF-32 TFBS which is a chromatin insulator and affects the regulation of gene transcription [85]. BEAF-32 is also connected to the development of photoreceptor differentiation during embryogenesis [88]. Therefore, these candidate TEs have the potential to affect the regulation of the flanking cold tolerant genes, as has been shown before for *D. melanogaster* TE insertions [84].

### TEs associated with inversions

TEs and other repetitive regions have been suggested to be involved in the origin of chromosomal inversions [118]. By studying three inversions recently characterized in *D. montana* across its distribution ([46] and Poikela et al., in prep), we were able to search for the presence of TEs near inversion breakpoints. These inversions were found on chromosomes X and 4, which also have the highest TE density. Perhaps not surprisingly, we found TEs nearby all the breakpoints. The breakpoints of the large chromosome 4 inversion that is fixed in Crested Butte harbors five TE insertions that are exclusive to Crested Butte and not found in the other populations. Similarly, the breakpoints of the small chromosome 4 inversion found in Oulanka and Kamchatka show TE insertions that are specific to these populations but are not found in the others. These findings could suggest that some of these TE insertions found around the inversion breakpoints may have acted as substrates for ectopic recombination and facilitated the origin of the inversions, although further analysis would be needed to confirm it [118]. Finally, the breakpoints of the X inversion have several TE insertions that are present in all *D. montana* populations, in accordance with the inversion being fixed in *D. montana*. Some of these insertions could have potentially played a role in the establishment of that X inversion ∼3 MY [46], but would require an in-depth investigation into the TEs between *D. montana* and other species of the *montana* phylad (*D. flavomontana, D. borealis, D. lacicola)*. Overall, these small insertions can greatly impact the formation of large rearrangements, potentially leading to significant evolutionary outcomes. For example, the large chromosome 4 inversion is associated with SNPs linked to cold/climate adaptation in *D. montana* and with a reduction in gene exchange between *D. montana* populations ([43] and Poikela et al., in prep.). Additionally, the X inversion is linked to the speciation process between *D. Montana* and *D. flavomontana* [46,119].

## Conclusions

Studying TEs in non-model species expands our understanding of TE dynamics throughout zootaxa, and their various impacts on the genome evolution. Adding carefully curated and a non-redundant TE library from non-model species to the public repositories significantly contributes to this mission. We recovered 555 curated non-redundant TE consensuses with remarkable numbers of new families and subfamilies comprising over 90% of the TE library in *D. montana* genome for the first time, using genetically distant intraspecies samples. TE contribution to the whole genome content is generally smaller in *D. montana* than its relative *Drosophila* species assuming that the differences in the quality of the genome assemblies between different species is not the case. As expected TEs were non-randomly distributed, we also found a higher density in gene-poor regions, and the X chromosome compared to autosomes. However, the TE abundance at the order level seems to vary across different species which worths more investigation from the evolutionary standpoint. Finally, we found several TEs around three inversion breakpoints for the first time in *D. montana* which are worth further studies on their potential impact on generating the inversion.

## Supporting information

D. montana TE library

## List of abbreviations

TE: transposable element
NA: North America
NE: North Europe
FE: Far East
TFBS: Transcription factor binding sites
LTR: long terminal repeats
LINE: long interspersed nuclear elements
TIR: terminal inverted repeats
RC: rolling circles

## Acknowledgements

The authors would like to thank Sara Lommi for the work in the laboratory and in the fly strain maintenance. Analyses were carried out using CSC (Finnish IT Center for Science) services.

## Availability of data and materials

The datasets supporting the conclusions of this article are included within the article and its additional files. The genome assemblies are retrievable with project name PRJNA828433, biosamples SAMN27782981-SAMN27782986. The TE library is provided as Additional file 4. The scripts developed for the purpose of this study are available in the Github repository, https://github.com/tahami-ms/TE-project

## Supplementary information

File name: Additional file 1

File formats: xls

Title of data: **Supplementary Table S1**. Sampling locations, years, and summary of the samples sequenced. **Supplementary Table S2.** Samples used for PacBio sequencing. **Supplementary Table S3.** Illlumina raw reads used for population studies. **Supplementary Table S4.** Manual curation criteria for de novo TE library construction in *D. montana*. **Supplementary Table S5.** Eu and heterochromatin coordinates for each chromosome in the five studied genomes. **Supplementary Table S6.** Number of genes transferred from the reference (monSE13F37) to other *D. montana* genomes. **Supplementary Table S7.** Selected cold-tolerant genes in *D. montana*. **Supplementary Table S8.** Transcription factor motifs used in this study that are related to stress. **Supplementary Table S9.** Reference genome coordinates for inversions found in *D. montana*. **Supplementary Table S10.** Statistics of identified TE consensus sequences using the REPET pipeline before and after manual curation. **Supplementary Table S11.** Description of the TE consensus sequences in the library of *D. montana*. **Supplementary Table S12.** Linear models testing for significant associations between raw reads, contigs and scaffold metrices. **Supplementary Table S13.** Eigenvalues from principal component analysis of SNPs. **Supplementary Table S14.** Distribution of TE insertions in different genomic regions. **Supplementary Table S15.** Population frequency of potentially active TE insertions. **Supplementary Table S16**. List of putatively adaptive TEs associated with cold-tolerant genes and present in the five genomes analyzed. **Supplementary Tables S17.** List of putatively adaptive TEs associated with cold-tolerant genes and present in 2-5 of the genomes analyzed. **Supplementary Table S18.** List of putatively adaptive TEs associated with cold-tolerant genes and present in only one of the genomes analyzed. **Supplementary Table S19.** TEs present in the five genomes analyzed that contain promoter sequences up-or downstream of cold-tolerant genes. **Supplementary Table S20.** Cold-tolerant related TFBS motifs found in TEs present in the five genomes and located up-/downstream or in the first intron of stress genes. **Supplementary Table S21.** Population frequency of the 34 TE insertions.

File name: Additional file 2

File formats: docx

Title of data: Supplementary method

File name: Additional file 3

File formats: docx

Title of data: **Supplementary Figure S1.** TE density per 5 kb bins of each of the five *D. montana* genomes analyzed, plotted by each chromosome. **Supplementary Figure S2.** Locations of fixed and polymorphic inversions with reference to centromeres.

File name: Additional file 4

File formats: txt

Title of data: TE library

## Authors’ contributions

MST: Wrote the manuscript, prepared the tables and figures, developed code and carried out the analyses, CVC: Contributed to analyses and proofreading. NP: Contributed to analyses, writing, and proofreading. MCZ: Contributed to analyses, developed code, proofreading. JG: Contributed to conceptualization, writing, and proofreading. MK: conceived and designed the project, contributed to writing, proofreading, and project management. All the authors read and approved the final manuscript.

## Ethics approval and consent to participate

Not applicable

## Consent for publication

Not applicable

