## Supplementary figures for "Transposable elements in a cold-tolerant fly species, *Drosophila montana*: a link to adaptation to the harsh cold environments"

**Supplementary Figure S1.** TE density per 5 kb bins of each of the five *D. montana* genomes analyzed, plotted by each chromosome. Chromosomes orders from top to down are: chr 2L, chr 2R, chr 3, 4, chr 5, and chr X.

| 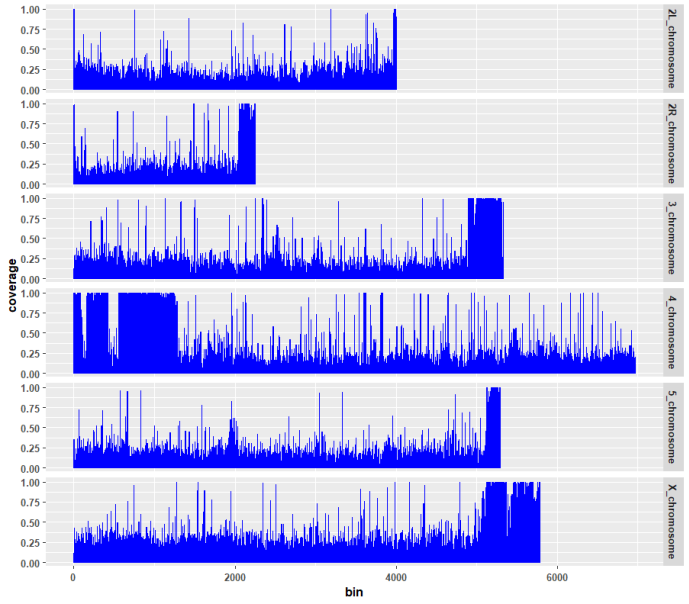 | 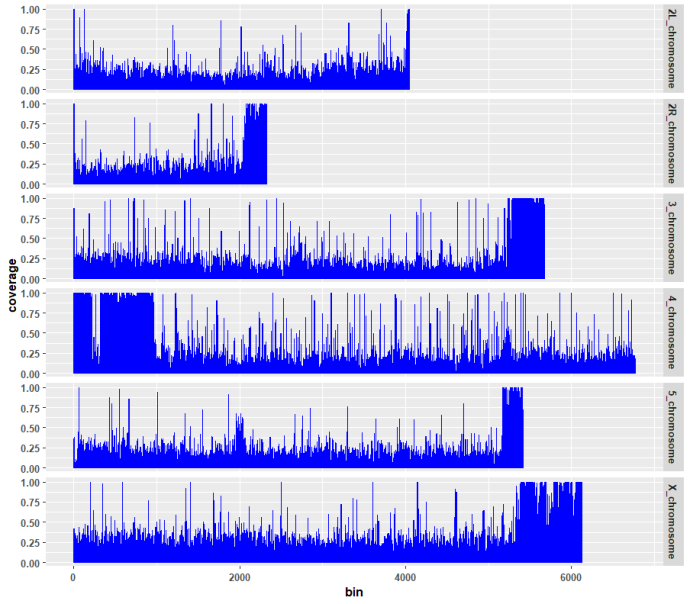  Jackson |
| --- | --- |
| 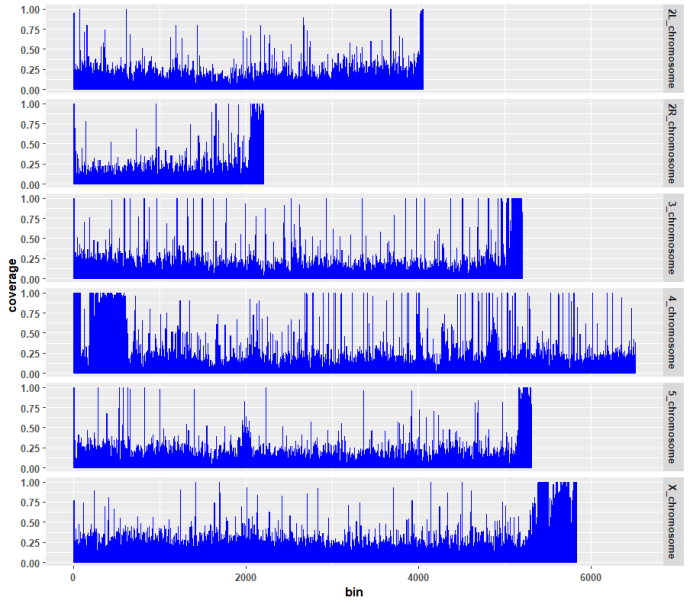  Seward | 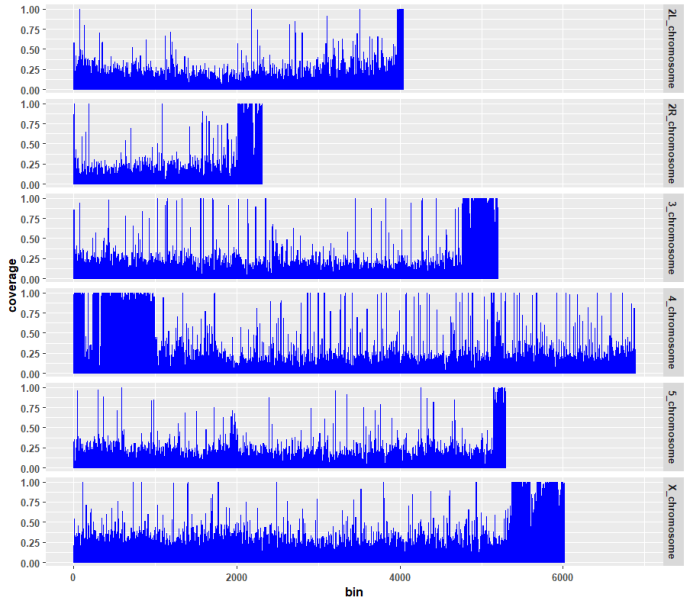  Kamchatka |
| 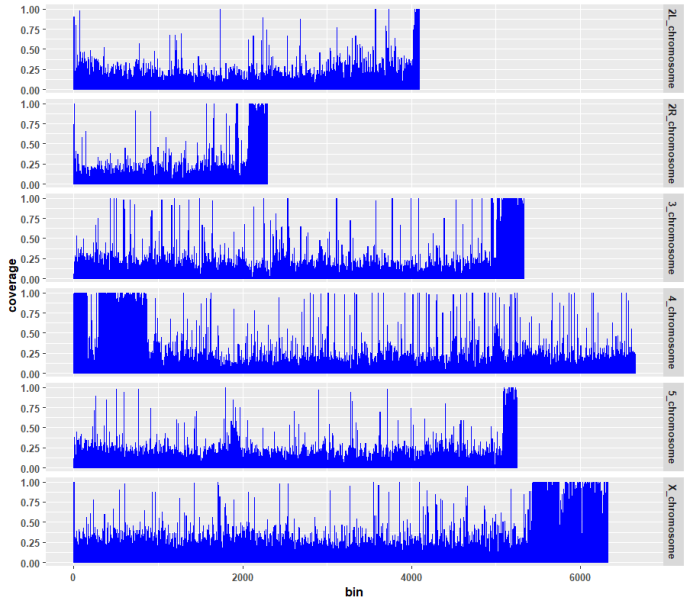 | Oulanka |

**Supplementary Figure S2.** Locations of fixed and polymorphic inversions with reference to centromeres. Proximal and distal breakpoints are not shown for the small polymorphic inversion.

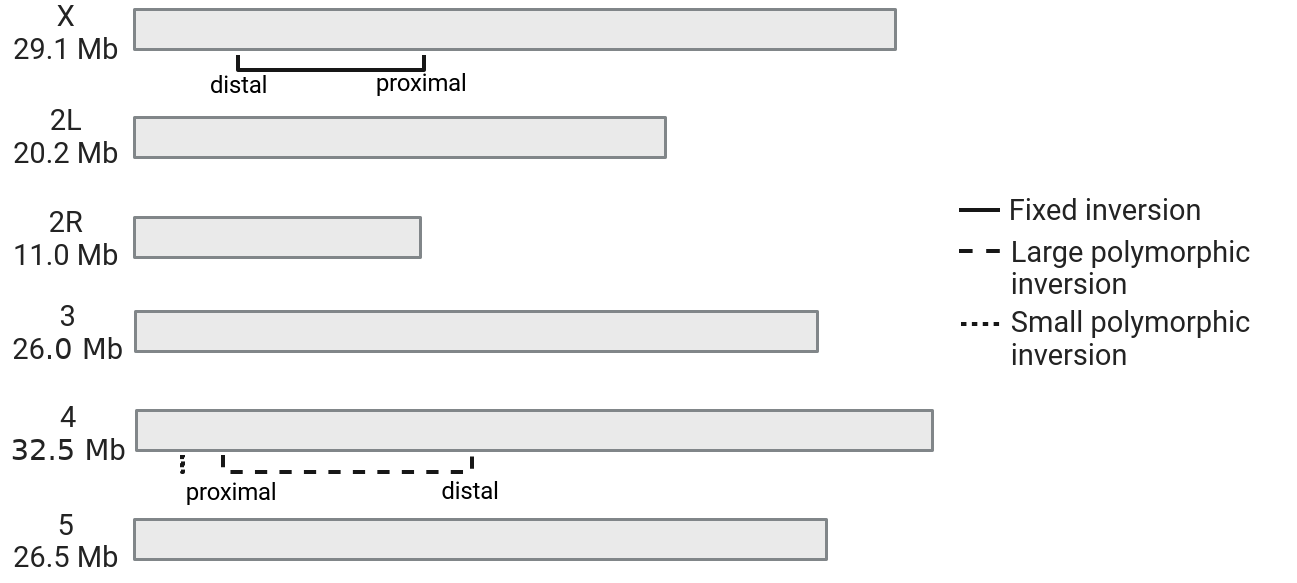
