## Supplementary method for "Transposable elements in a cold-tolerant fly species, *Drosophila montana*: a link to adaptation to the harsh cold environments"

**Supplementary Material**

**Manual curation guideline**

We defined a set of conditions to reduce the number of artifacts and redundant TE sequences (Table S3). For autonomous TEs, we accepted the TE consensus if it fulfilled the criteria of having at least one full length copy throughout the genome. For non-autonomous TEs, we accepted the consensus if there were more than one copy in the genome. We revised/complemented our curation by masking TE consensus on the updated internal TE library implemented in CENSOR [1,2].

After the manual curation, consensus sequences were classified at the family level following the comparison with the *Repbase* library for eukaryotes [3], *Flybase* [4] and *Dfam* [5,6] libraries for *Drosophila* using *CD-HIT* web interface [7], module “cd-hit-est-2d” with -c 0.8, -aS 0.8 and -r ‘yes’ parameters (Accessed Oct 2021). Consensus sequences that were not clustered with a known TE based on the 80-80-80 rule, were masked to the reference library in CENSOR for classification based on their match. Chimeric consensuses were trimmed using Geneious Prime 2021.2.2 [8]. After classification, all five libraries were combined and clustered together using *CD-HIT* web interface, module “cd-hit-est” with -c 0.8 -aS 0.8 and -r ‘yes’ parameter sets (Accessed Oct 2021). to remove redundancies and the longer TE from each cluster was kept. If several TE consensuses met the 80-80-80 rule against one known TE family but were not clustered together, we gave them a subfamily rank.

We ran *MCHelper* (beta version 0.8.0) with each of the five libraries to assess their quality [9]. MCHelper reduces library redundancy, extends the consensus sequences, if needed, filters out false positive sequences, including satellites, multi-copy genes, and rRNA sequences, and performs structural checking based on the consensus sequence classification. Our MCHelper analysis validated all consensus sequences except three that after visually inspecting MSA and TE+aid plots, were identified as chimeric sequences and were thus discarded.
